# Insect galls thrive on salt stressed big sagebrush host

**DOI:** 10.1101/821140

**Authors:** Cindy T Wu

**Affiliations:** Jelly

## Abstract

The sagebrush steppe is estimated to occupy 4.4 percent of the total land area in the United States. In this ecosystem, the most common sagebrush species in most areas is big sagebrush, *Artemisia tridentata*. Big sagebrush provide a common host for galling insects. We observed that even in salt stressed environments big sagebrush grow and galling insects use them as hosts. In this study we test the hypothesis that there is a significant difference in insect gall density on salt stressed big sagebrush compared to unstressed big sagebrush. To test this, we collected data from four salt stressed sage plots and four unstressed plots at Columbia National Wildlife Refuge in Othello, Washington during 2010. We report significantly greater average gall density on plants in salt stressed plots. We collected four types of galls (artichoke, white fuzzy, black fuzzy, smooth, and other) and observed no significant difference in the diversity of galls on sage in salt stressed plots compared to the control. Further studies are needed to understand why salt stressed sage show greater density of galls, why insect galls seem to thrive on salt stressed sage.

## 1 Introduction

In the sagebrush steppe area, big sagebrush (*Artemisia tridentata*) provide a common host for galling insects. The sagebrush steppe occupy an estimated 108 million acres of the United States or 4.4 percent of the total land area in the country[1]. Little is known about the sage-associated insects in these cold-desert regions[2][3]. Livestock grazing, the introduction of invasive species and other land uses greatly affects the sagebrush-steppe ecosystem[2]. Despite the large spatial extent and economic importance of sagebrush, the insects associated with sage are largely left unexplored[3].

Previous observations that water stress and grazing stress lead to fewer insect galls on the big sagebrush (B. Chowdhury, L. Thomason, personal communication, June 4, 2010) inspired my exploration of how salt stress affects gall density in big sagebrush. Comparing gall count between salt stressed and salt unstressed big sagebrush systems allow for an indirect way to assess how excess amounts of salt affect big sagebrush gall interactions.

Big sagebrush intermingles to a limited extent with strict halophytes, plants that tolerate or even demand sodium chloride, including the halophyte black greaswood or *Sarcobatus vermiculatus* [4]. Big sagebrush germinates well at an osmotic pressure of −0.253 mPa or lower[5]. Black greasewood which thrives at an osmotic potential of −4.2 mPa provides a system in the sagebrush steppe for us to identify salt stressed environments[6]. Because plant fitness changes are governed by biogeochemical processes occurring during soil development and plant succession, we use black greasewood as an indicator of salinity. Using presence of black greasewood as an indicator of salt stress, we establish an indirect way to evaluate the consequence of salt stress on insect gall formation in big sagebrush.

## 2 Methods

We collected data in the Columbia National Wildlife Refuge in Othello, Washington (46° 54’ 49” N, 119° 14’ 14” W) during the second and fourth weekend of May 2010. We used presence of black greasewood to indicate salt stressed soils.

We surveyed 100 square meter circle plots by measuring a 5.64 meter radius

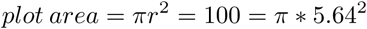

of sage-steppe with at least one black greasewood plant in salt stressed plots and no black greasewood in the control plots. To control for water stress none of our plots contain golden currant which requires excess water that big sagebrush generally avoids[7].

In each plot, we measured the height, length and width of each big sagebrush brush in centimeters (height: longest measurement from the ground, length: longest measurement across the plant, width: measurement of plant perpendicular to the length using the center of the length as the intersection between length and width). We removed all galls on each sage plant in the plot and characterized the gall types. We separated gall types into the following categories: artichoke, white fuzzy, black fuzzy, smooth, and other. Artichoke are green smooth galls with overlapping appendages, white fuzzy are galls with white fur-like coatings, black fuzzy are galls with black coarse fibers, smooth are galls with green-red smooth outer coatings, and other are galls that do not meet any of the above criteria.

I calculated the average gall density per cubic meter by dividing the number of galls per bush over the estimated volume of the bush in cubic meters. I estimated the volume of each bush by averaging the spherical volume and rectangular prism volume of each plant

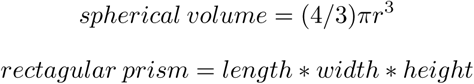

I averaged the gall densities for all bushes in each plot. I took the mean value of the mean gall density for each plot to report pooled mean gall density per cubic meter in each plot. I used a two-sample assuming unequal variances t-test to compare the two different treatments: salt stressed and control.

I calculated the average number of each gall category (artichoke, white fuzzy, black fuzzy, smooth and other) per bush in the observed plot. I used a two-factor ANOVA to compare the gall diversity of sagebrush in salt stressed plots to that of sagebrush in control plots. I used the two-factor ANOVA to evaluate the evenness in gall category distribution.

## 3 Results

Big sagebrush plants in plots of salt stressed soils had higher average gall density (M = 409.71, SE = 90.7) than those in the control plots (M = 80.59, SE = 18.0), t(3) = 3.5, p < 0.05 (Fig. 1). We collected data from four salt stressed plots and four control plots. Salt stressed plots and control plots show no significant difference in gall category diversity. However, the gall category fauna shows an unequal distribution of galls per bush (Two-Factor ANOVA: n = 20, *α* = 0.05, gall type: F = 212.13, F-critical = 2.71, df = 4, p < 0.01; stress condition: F = 0.96, F-critical = 2.36, df = 7, p > 0.01; Fig. 2.) We report no difference in the galls of the same category collected from salt stressed and control plots. Of the collected galls, one artichoke gall hatched out an insect of the order Diptera.

**Figure 1:**
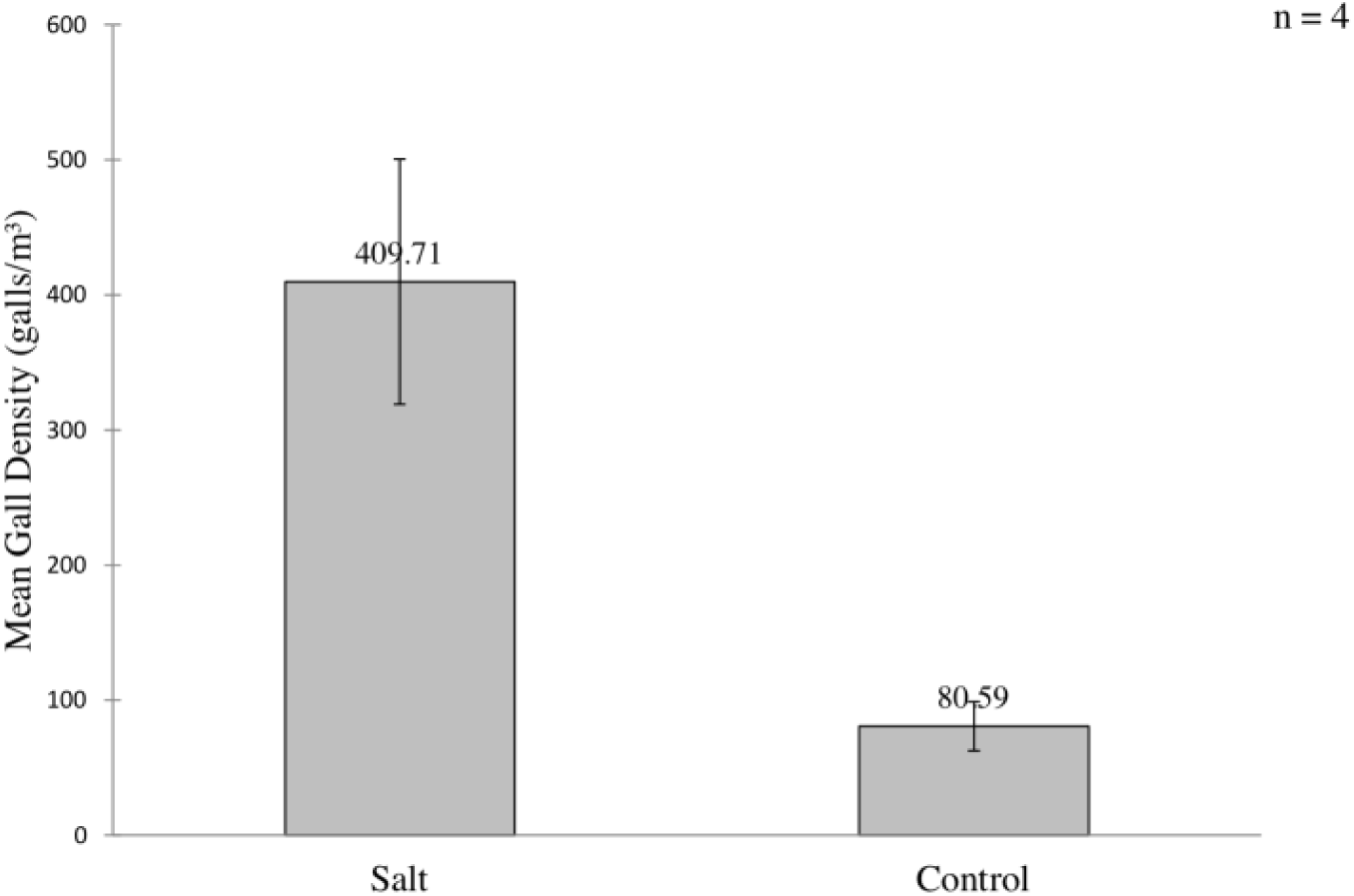
A. Tridentata shows greater gall density in regions of salt stressed soils. (Two-Sample Assuming Unequal Variances onetail t-Test: n = 4, *α* = 0.05, df = 3, t = 3.5, t-critical = 2.35, p < 0.05)

**Figure 2:**
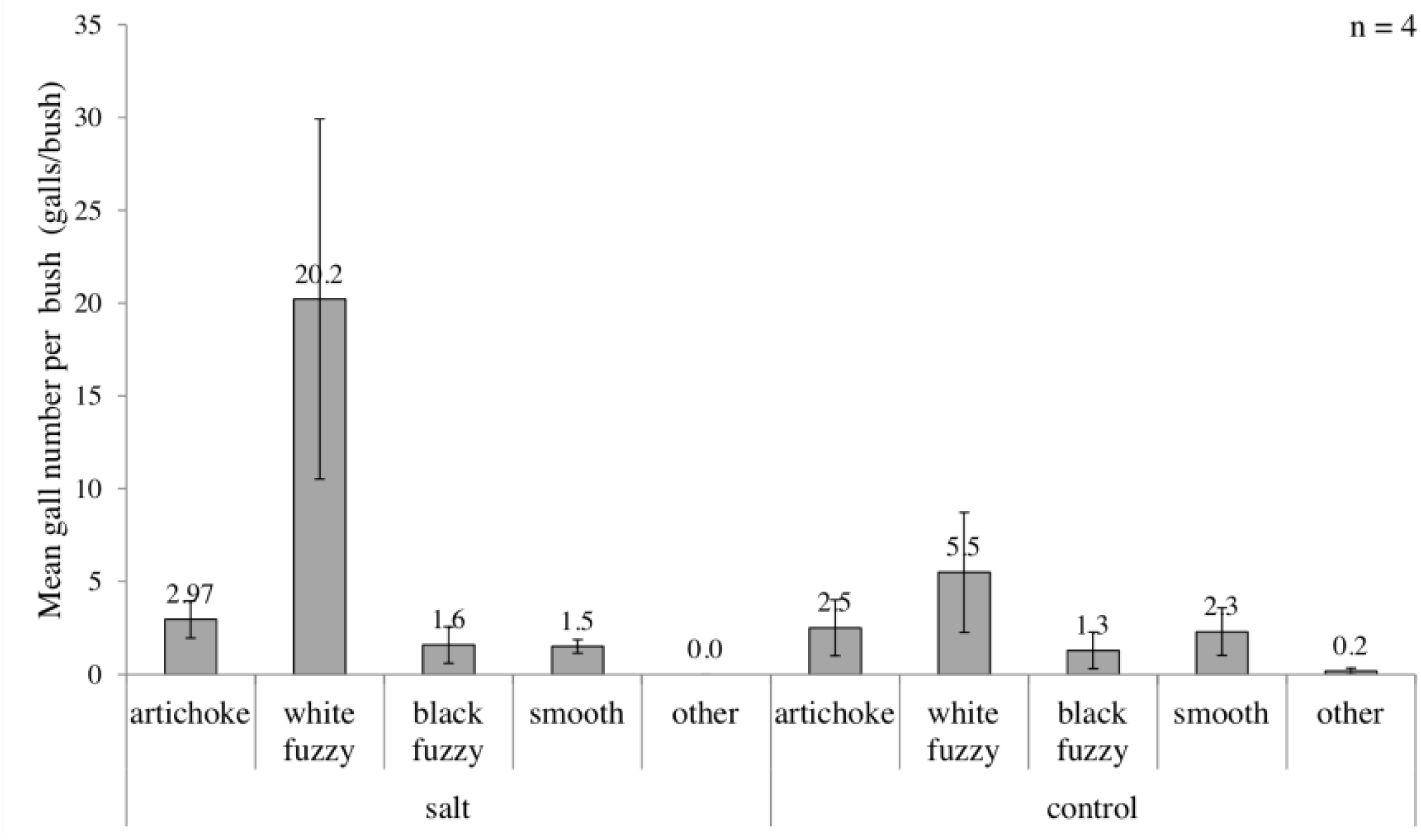
Unequal gall diversity does not differ between A. Tridentata in salt stressed compared to control conditions. (Two-Factor ANOVA: n = 20, *α* = 0.05, gall type: F = 212.13, F-critical = 2.71, df = 4, p < 0.01; stress condition: F = 0.96, F-critical = 2.36, df = 7, p > 0.01)

## 4 Discussion

Because not all stresses are created equal, the salt stress may upregulate growth hormones in certain parts of big sagebrush encouraging gall formation. Recent research shows that *Arabidopsis thaliana* under mid salt stress promotes auxin accumulation in developing primordia[8]. Plant development and growth can be stimulated by salt stress. Because the active differentiation of plant growth and tissue is largely controlled by the galler insect genes, if the insect can secrete bioactive molecules that differentiate meristematic tissue salt stress may induce growth in the gall areas. I further examined the relative bush size differences between studied conditions and found that there is no significant difference between the sage plant sizes (Two-Sample Assuming Unequal Variances one-tail t-Test: n = 4, *α* = 0.05, df = 3, t =-1.65, t-critical = 2.35, p > 0.05). The analysis of size difference shows that greater gall density on salt stressed big sagebrush is not attributable to a difference in plant size.

The phenomena of galling insects using big sagebrush as host involves multiple individual chemical pathways. The reason for the reported greater gall density on sagebrush plots in salt stressed soils cannot be determined by this study. The insect gall behavior on big sagebrush can be a good system for indicating of the consequences of a stressor such as salt, water, or grazing. However, with the gall model the pathways associated with different stress variables remain unclear. The reported observations of (1) greater density of galls on salt stressed big sagebrush, (2) diverse gall insect population inhabiting big sagebrush, and (3) an insignificance between gall diversity in the salt stressed and not salt stressed plots that these observations are still unexplained. It is worth noting that we do not yet know what species of insects are creating these galls. Further study is necessary to understand how and why these galls exhibit these behaviors on big sagebrush.

## 5 Acknowledgements

I thank P. Dee Boersma, Amy Van Buren, Bashira Chowdhury, Jessica Jang, Estella Leopold, Denny Luan and Jaehyung Park for helpful discussions, Richard Chong, Bashira Chowdhury and Lindsay Thomason for help with field work and data collection, and Cecilia Nem and Jessica Jang for constructive comments on the manuscript. Any mistakes in this manuscript are my own.

